# Multi-level, multi-body atomic interaction graphs for machine learning-based prediction of protein-ligand binding energies

**DOI:** 10.64898/2026.06.05.730001

**Authors:** Tram TH. Le, Huy Vo, Nhat HM. Nguyen, Binh T. Nguyen, Duc Duy Nguyen

## Abstract

Accurate prediction of binding affinity is crucial for rational drug design and discovery. Traditional computational methods often rely on complex scoring functions that incorporate a multitude of physical and chemical descriptors, leading to high computational demands and sometimes limited generalizability. In this work, we propose a novel scoring function that models multi-level, multi-body atomic interactions using graph-based representations. Our method constructs comprehensive interaction graphs that incorporate both pairwise and triplet-wise atomic features that help capture cooperative spatial patterns essential for binding affinity prediction. By employing a feature fusion strategy, GMI-Score maintains model simplicity while enhancing accuracy. Extensive evaluation across multiple datasets, such as PDBbind v2013, PDBbind v2016, PDBbind v2020, CSAR-NRC-HiQ, and PDBbind-Redocked, demonstrates that our model consistently outperforms state-of-the-art scoring functions, achieving Pearson correlation coefficients up to 0.877. Furthermore, it retains strong predictive power under strict data leakage controls and realistic docking conditions to highlight its robustness and generalizability.

**Scientific Contribution:** In this study, we present a scoring methodology that systematically captures higher-order atomic interactions within a unified graph framework, making a conceptual shift in cheminformatics scoring functions. Its consistent outperformances of existing methods and strong validity under redocked and withheld atascenarios demonstrate its utility for broad-scale molecular modeling applications and open heminformaticsworkflows.

## 1 Introduction

Protein-ligand interactions are fundamental to nearly all biological processes, such as enzyme catalysis,signal transduction, and molecular recognition [3, 5, 16, 67]. These interactions typically depend on non-covalent forces, including hydrogen bonds, ionic interactions, hydrophobic effects, and van der Waals forces. These interactions provide critical insights into systems biology, revealing mechanisms such as signaling pathways [8], allosteric regulation [9], and feedback control that underlie complex cellular processes [17].

Given their central role in cellular function, quantifying protein-ligand interactions becomes essential, particularly in fields like drug discovery, where precise control over these interactions can determine therapeutic efficacy. The strength of these interactions is typically expressed through binding affinity, often quantified by the equilibrium dissociation constant K_d_ [2]. A lower K_d_ value indicates a higher affinity between the protein and ligand and vice versa. Accurate binding affinity predictions can be utilized in virtual screening, lead optimization, and cost-effective *de novo* drug design [53]. Consequently, developing robust computational methods to predict binding affinities has become a key objective in modern pharmacological research [57].

Traditionally, experimental methods such as isothermal titration calorimetry (ITC) or surface plasmon resonance (SPR) are used to determine protein-ligand binding affinity [12, 46]. However, these methods are usually expensive and time-consuming, making them unsuitable for large-scale drug screening [71]. As a result, there is a need for computational models that can effectively and accurately predict binding.In response, recent technological advances have led to the development of a variety of computational approaches for studying protein-ligand interactions. These include structure-based methods such as molecular docking, standalone scoring functions designed to predict binding affinity, and physics-based techniques like molecular dynamics simulations that analyze the stability and conformational dynamics of protein–ligand complexes [1].

In computational drug design, scoring functions are mathematical models used to estimate the protein-ligand binding affinity. They are commonly categorized into four types: physics-based, empirical, knowledge-based, and machine learning-based. Physics-based scoring functions focus on intermolecular interactions, such as van der Waals forces and electrostatic attractions [21, 30]. Empirical scoring functions use energy terms derived from molecular interactions and some empirical chemistry factors like hydrophobicity and metal-ligand interactions [47, 49]. Knowledge-based scoring functions rely on statistical analysis of known protein-ligand complexes to identify favorable interaction patterns [60]. With the growth of data, machine learning-based functions are gaining much more attention. These methods learn patterns from large training data and have been shown to outperform traditional methods in binding affinity prediction [4, 22, 29, 44, 51, 61, 74]. Despite extensive research throughout the years, protein-ligand interactions are still far from fully understood. As the binding processes are influenced by many factors ranging from spatial arrangement to interactions between the molecules, feature selection remains a difficult task in machine learning-based methods [66]. Many existing works focus on the interactions between atom pairs in the protein-ligand complex. RF-score utilizes a feature vector of occurrence-specific atom type pairs within a cutoff distance [4]. ECIF extends this by considering atomic connectivity, valence, and hybridization [51]. At a more detailed level, GGL-ETA-Score constructs subgraphs based on atom types and computes features through summation of the generalized Lorentz or Exponential function over atom pairs [50]. A similar approach by Fangqiang Zhu et al. also uses feature vectors consisting of the summation of a neural network over atom pairs [74]. However, we hypothesize that pairwise interactions may not fully capture the characteristics of binding processes. We believe that multi-level interactions can provide a better representation for the protein-ligand complex and enhance the performance of machine learning-based scoring function models.

Models such as DimeNet [19], GemNet [20], and PaiNN [52] incorporate angular and dihedral information over bonded molecular graphs to improve expressiveness. However, these models were designed and benchmarked primarily on covalently bonded small-molecule systems, where connectivity is chemically prescribed and training datasets are relatively modest in scale. In contrast, the protein-ligand scoring problem presents a fundamentally different regime: interaction graphs are defined by spatial distance cutoffs rather than covalent bonds, labeled data is available at large scale, and inference efficiency is critical for practical screening workflows. Direct application of these architectures is therefore computationally prohibitive, as models such as GemNet-OC require quadratic or cubic operations through multi-hop directional message passing and process orders of magnitude fewer samples per GPU-second than simpler models at inference time.

In this work, we present GMI-Score (Graph geometric learning with Multi-body Interactions based scoring functions), a novel framework for modeling protein-ligand binding affinity. While prior models such as RI-Score and GGL-Score [44, 50] rely on graphs constructed from pairwise atomic interactions, GMI-Score introduces a more expressive representation by constructing multi-body interaction graphs that capture higher-order geometric motifs, including triplet-wise interactions. In addition to expanding the graph structure, GMI-Score incorporates a multi-level interaction fusion mechanism that jointly learns from different interaction orders. This design enables the model to capture cooperative spatial patterns that are often overlooked by pairwise methods. By learning from both fine-grained and collective structural features, GMI-Score provides a more faithful representation of the binding interface. This design consciously trades the full expressive power of equivariant architectures for practical scalability, providing the geometric inductive bias necessary to surpass pairwise methods without incurring the computational overhead that makes geometric GNNs impractical at protein-ligand screening scale.

Evaluated across multiple standard and challenging benchmarks, including CASF-2013 [37], CASF-2016 [59], PDBbind v2020, CSAR-NRC-HiQ [13], and PDBbind-Redocked [54], our method cachieves better results than other state-of-the-art models. The strong performance and efficient design of GMI-Score underscore its potential as a powerful tool for accelerating drug discovery in both academic and industrial settings.

## 2 Results and Discussion

In this section, we present the scoring power of our proposed multi-level model, GMI-Score, for the benchmark datasets.

### 2.1 Datasets

To evaluate the predictive power of our proposed method, we utilize the PDBbind v2013, the PDBbind v2016, and the PDBbind v2020 datasets, which are standard datasets for the binding affinity prediction task. The PDBbind datasets are divided into three nested subsets: the general set, the refined set, and the core set, with each subset being contained within the previous one. The general and refined sets, after excluding the core set, are used as training data for our model, whereas the core set is used as the testing set. Despite being a well-acknowledged dataset, several studies have demonstrated concerns regarding the reliability of the PDBbind datasets as a benchmark for scoring functions. The protein-ligand complexes in the subsets are not consistent in terms of quality. Specifically, the core set contains protein-ligand complexes with higher resolution and binding affinity compared to complexes in the general and refined sets [38].

In order to provide a more comprehensive assessment of our method’s predictive ability, we conducted further evaluations on two external test sets: the CSAR-NRC-HiQ and PDBbind-Redocked datasets.

The CSAR-NRC-HiQ dataset [13] is a valuable resource for validating molecular docking and scoring models. It originally contains a total of 343 high-quality protein-ligand complexes, which are divided into two distinct subsets of 167 and 176 complexes.

To create an independent test set, we combined both subsets and then removed any complexes that overlapped with the PDBbind database. This filtering process resulted in a final, non-overlapping set of 49 complexes, which we used for external validation of our model’s performance.

The PDBbind-Redocked [54] is used for evaluating a scoring function’s ability to perform in real-world scenarios, where co-crystal structures aren’t available. Instead of using the experimentally determined poses from the PDBbind v2016 dataset, this dataset uses redocked structures generated by Surflex-Dock [24]. To assess a scoring function’s performance on this dataset, all complexes that are part of the CASF-2016 set are used as a test set.

All the datasets are shown in Table 1.

**Table 1:** Summary datasets used in our work.

| Datasets | Training sets |  | Test set |
| --- | --- | --- | --- |
|  | General set | Refined set |  |
| PDBbind v2013 | - | - | 195 |
| PDBbind v2016 | 12998 | 3772 | 285 |
| PDBbind v2020 | 23496 | 5316 | 285 |
| CSAR-NRC-HiQ | - | - | 49 |
| PDBbind-Redocked | - | - | 285 |

### 2.2 Hyperparameters optimization

To achieve the optimal hyperparameters for the binding affinity prediction task, we followed the hyperparameter settings from our previous work [50] for the pairwise interaction model, i.e., GMI-Score-2. We then optimized the GMI-Score-3 scoring function by evaluating its performance across a range of hyperparameters (*t, k*) and cutoff distances (*c*). We also hyperparameter-tune the model with grid search to identify optimal hyperparameters for our chosen machine learning models, aiming to maximize the correlation between predicted and experimental binding affinities. To balance thoroughness with computational feasibility, we performed GridSearchCV on the first training dataset (PDBBind v2016) to identify optimal hyperparameter configurations, and then applied these same configurations across all other training-test scenarios. This approach follows established practice in machine learning, where hyperparameters optimized on one representative dataset are transferred to related tasks to avoid overfitting to any specific dataset and to reduce computational overhead.The resulting best-performing models for each dataset served as the foundation for further analysis. The performances of various Gradient Boosting models on the CASF-2016 [59] benchmark trained on general set are presented in Table 2 and Table A1.

**Table 2:** Hyperparameter selection for the Gradient Boosting model of GMI-Score-3 on the CASF-2016dataset. RMSE is in kcal/mol units.

| Model | Train with Refined |  | Train with General |  |
| --- | --- | --- | --- | --- |
| | $R_p$ | $RMSE$ | $R_p$ | $RMSE$ |
| Model 1 | $0.836 \pm 0.002$ | $1.742 \pm 0.004$ | $0.860 \pm 0.002$ | $1.638 \pm 0.007$ |
| Model 2 | $0.836 \pm 0.003$ | $1.703 \pm 0.003$ | $0.857 \pm 0.002$ | $1.623 \pm 0.007$ |
| Model 3 | $0.824 \pm 0.006$ | $1.734 \pm 0.021$ | $0.850 \pm 0.006$ | $1.630 \pm 0.025$ |
| Model 4 | $0.836 \pm 0.002$ | $1.709 \pm 0.007$ | $0.857 \pm 0.001$ | $1.626 \pm 0.002$ |

To explore the potential for improved predictive power, we investigated the fusion of features derived from GMI-Score-3 and GMI-Score-2. Integrated feature sets were 5constructed by combining these two scoring functions at various ratios. For each benchmark dataset and each feature ratio, the optimized machine learning model was trained and evaluated with the five independent runs using various random seeds to ensure robust performance assessment.

### 3.2 PDBbind v2016 Benchmark

Our findings indicate that the GMI-Score-2, through its atom-specific feature representation, and the GMI-Score-3, with its atom-wise approach, capture different facets of the complex binding affinity between proteins and ligands. While each scoring function possesses inherent strengths, our analysis suggested that not all constituent features contribute equally to the predictive accuracy; indeed, some may have a negligible or even detrimental impact. This observation motivated the development of a fusion strategy focused on integrating the most relevant variables from both scoring paradigms. The primary objectives of this fusion were twofold: firstly, to achieve a more lightweight model by reducing the total number of input features, which in turn lessens computational cost and can enhance model interpretability; and secondly, to elevate the predictive performance beyond what is achievable by either scoring function in isolation.

Utilizing the PDBbind v2016 dataset for evaluation, models were trained on a combination of their general sets, and their baseline performances were subsequently tested on the core set. We first established baseline performances for the individual scoring functions, obtaining Pearson correlation coefficients of 0.870±0.001 for the GMI-score-2 and 0.863 ± 0.002 for the GMI-Score-3 (Table 3). Specifically, the optimized fusion model achieved a Pearson correlation coefficient of 0.877 ± 0.001.

**Table 3:**
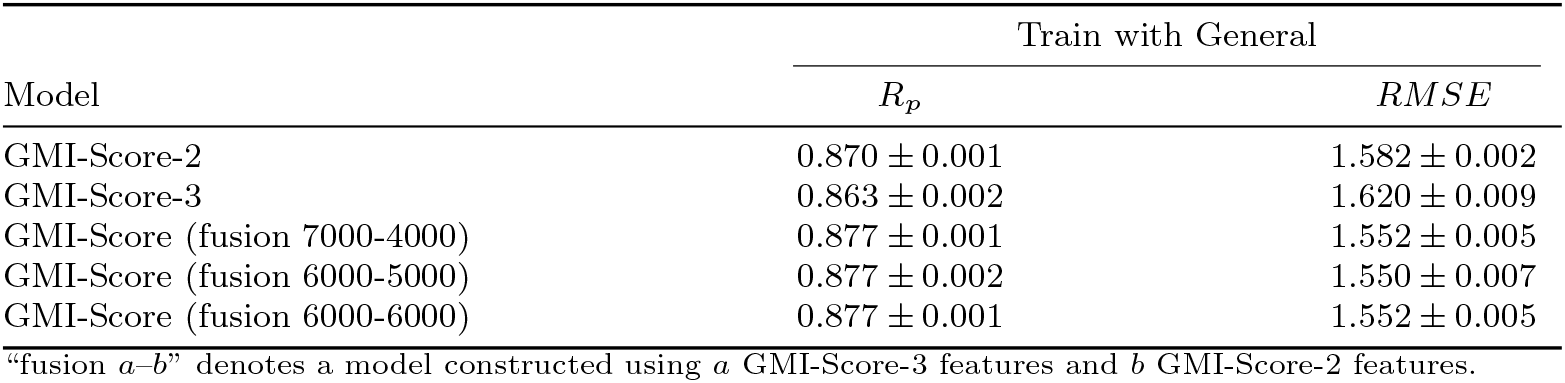
Performance comparison of individual (GMI-Score-2 and GMI-Score-3) and fusion models on the CASF-2016 dataset. The table reports the Pearson correlation coefficient (*R*_*p*_) and RMSE in kcal/mol.

**Table 4:**
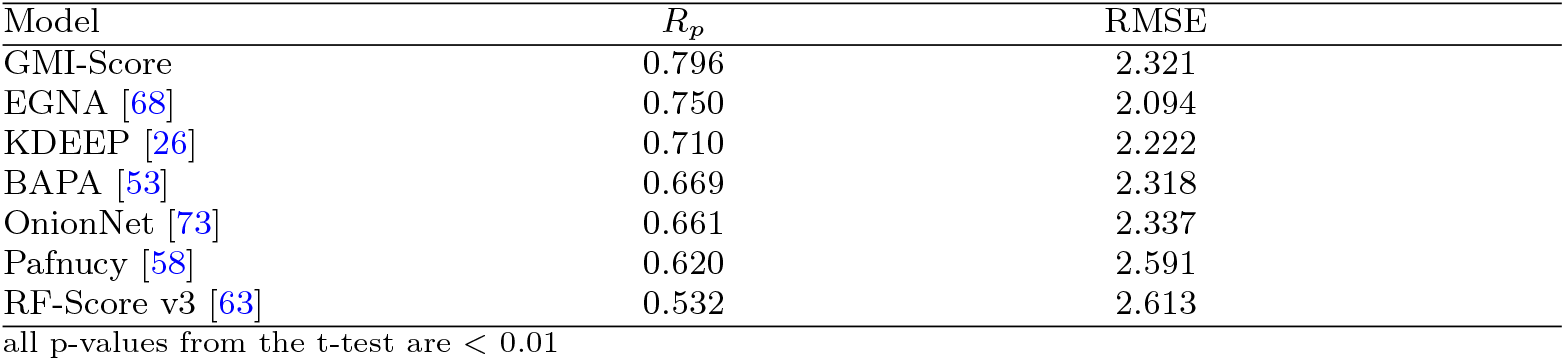
Pearson correlation with predictions on CSAR-NRC-HiQ dataset. RMSE is in kcal/mol units. The performances of other models are taken from [68].

Paired t-test have been applied to see if the difference of nearly 0.02 of *R*_*p*_ meaningful between GMI-Score and the two individual models (Table A2). The outcomes of these experiments were encouraging and demonstrated that the resulting hybrid models statistically significantly better than both individual models. This enhanced performance, together with the advantages of feature reduction, validates the effectiveness of our feature fusion approach. The fusion model GMI-Score (fusion 7000–4000) is used for all evaluations; hereafter, it is referred to as GMI-Score.

When benchmarked against other state-of-the-art models on the CASF v2016 dataset, our proposed GMI-Score performed competitively (Figure 1). We believe this improvement may be attributed to the synergistic combination of the GMI-Score-2’s capacity to capture overarching protein-ligand relationships and the GMI-Score-3’s detailed atom-wise insights. Our approach attempts to distill these strengths into a concise and computationally efficient feature set. Importantly, this fusion strategy appears to do more than simply preserve the predictive power of the individual components; it also seems to help mitigate their respective limitations. As a result, the proposed method is more robust and accurate for determining protein-ligand binding affinity than previous ones [4, 36, 50].

**Fig. 1:**
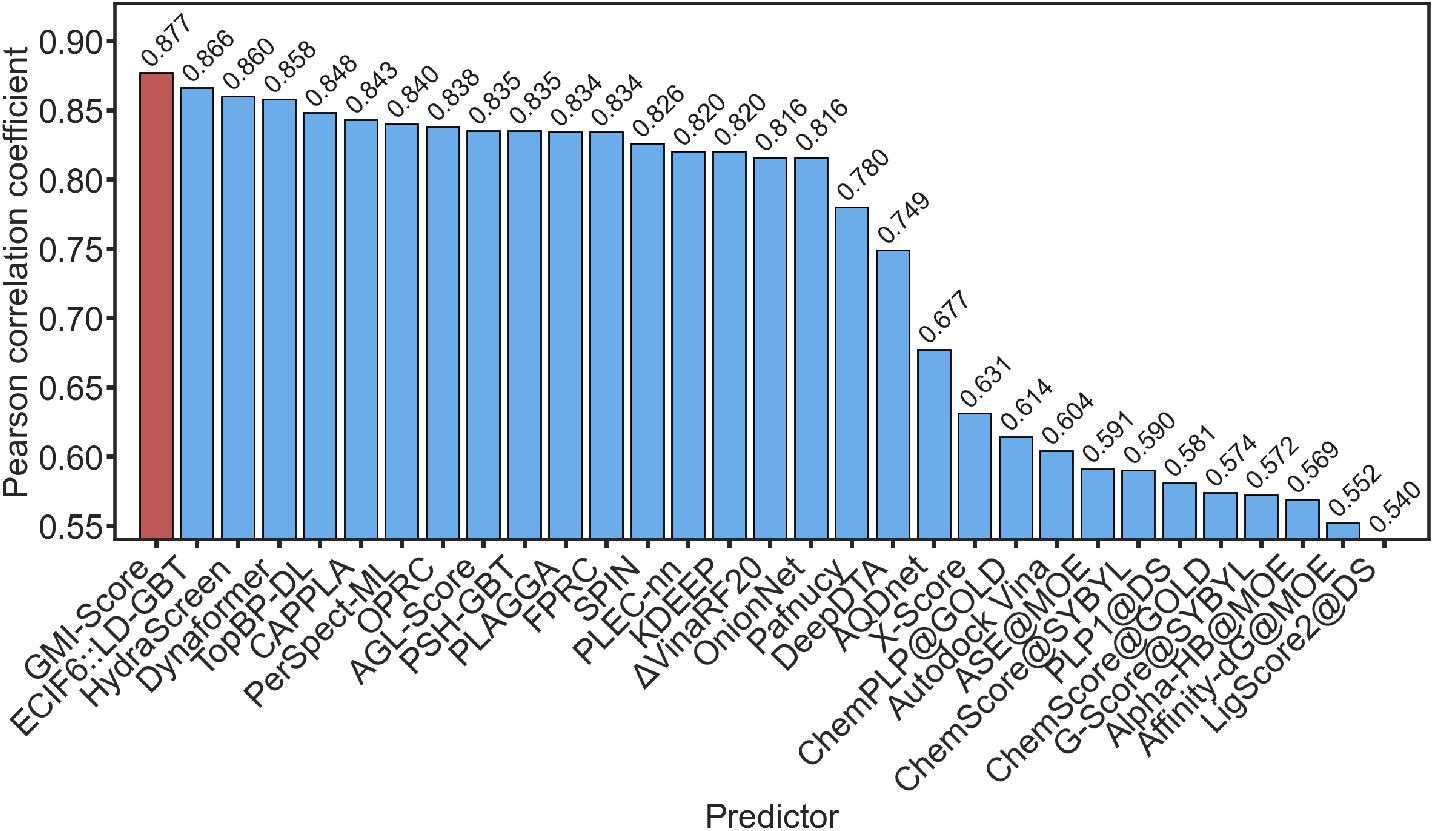
Comparison of Pearson’s correlation scores of various scoring functions on the CASF-2016 benchmarks [11, 27, 33, 39, 48, 50, 56]. Our proposed *GMI-Score* is shown in red, while other methods are shown in blue.

### 2.4 PDBbind v2020 Benchmark

We extended its application to the PDBbind v2020 dataset in order to validate the generalizability of ourGMI-Score. This dataset is a significantly expanded version of PDBbind v2016, encompassing 19,443 protein-ligand complexes. Notably, it retains the same core set as PDBbind v2016, facilitating a direct comparison of performance and allowing us to assess the model’s ability to handle a larger and more diverse dataset. Adhering to the established PDBbind evaluation protocol, we trained the GMI-Score framework on general and refined sets of PDBbind v2020. To mitigate computational costs associated with extensive parameter tuning on this larger dataset, we transferred the optimized parameters obtained from our experiments with PDB-bind v2016. This approach assumes a degree of parameter eneralizability across the two datasets, which is reasonable given the overlap in their core sets and the underlying physicochemical principles governing protein-ligand binding. Although fine-tuning the GMI-Score model on the v2020 dataset could potentially yield better performance, we chose to omit this step to highlight the feasibility and transferability of our method.

The trained GMI-Score was then tested on the PDBbind v2020 core set. As reported in Table A3, our method achieved a Pearson correlation coefficient (*R*_*p*_) of 0.872 ± 0.001. A strong linear correlation between predicted and experimental binding energies is further illustrated in the right panel of Fig. B1, which shows that most data points lie close to the diagonal line with RMSE=1.555 kcal/mol. The p-value of the t-test between the individual model and GMI-Score showed significance for both models trained with the refined and general sets (Table B4). This result is particularly noteworthy for several reasons. First, the high correlation coefficient indicates a strong linear relationship between the GMI-Score predictions and the experimentally determined binding affinities, demonstrating the model’s predictive accuracy. Second, when compared to the performance of separate scoring functions with fusion model on the same benchmark (0.862 ± 0.002 and 0.863 ± 0.002, respectively), this result underscores the generalization capability of our GMI-Score. The ability to maintain a high level of accuracy on a significantly larger and more diverse dataset, without extensive re-parameterization, suggests that GMI-Score has effectively captured the fundamental determinants of protein-ligand binding affinity. This enhanced performance is a critical advantage for practical applications in drug discovery, where models are often applied to novel protein-ligand systems not seen during training.

Further comparison with existing scoring functions is shown in Fig. 2, where GMI-Score achieves the highest Pearson correlation coefficient among all evaluated models. DeepDTA [45] and SPIN [11] show lower correlations (0.860 and 0.826, respectively), which demonstrates that GMI-Score provides better predictive accuracy and robustness on the PDBbind v2020 dataset over other state-of-the-art models. This pretrained model on this dataset is used for further evaluation on external test datasets and community use.

**Fig. 2:**
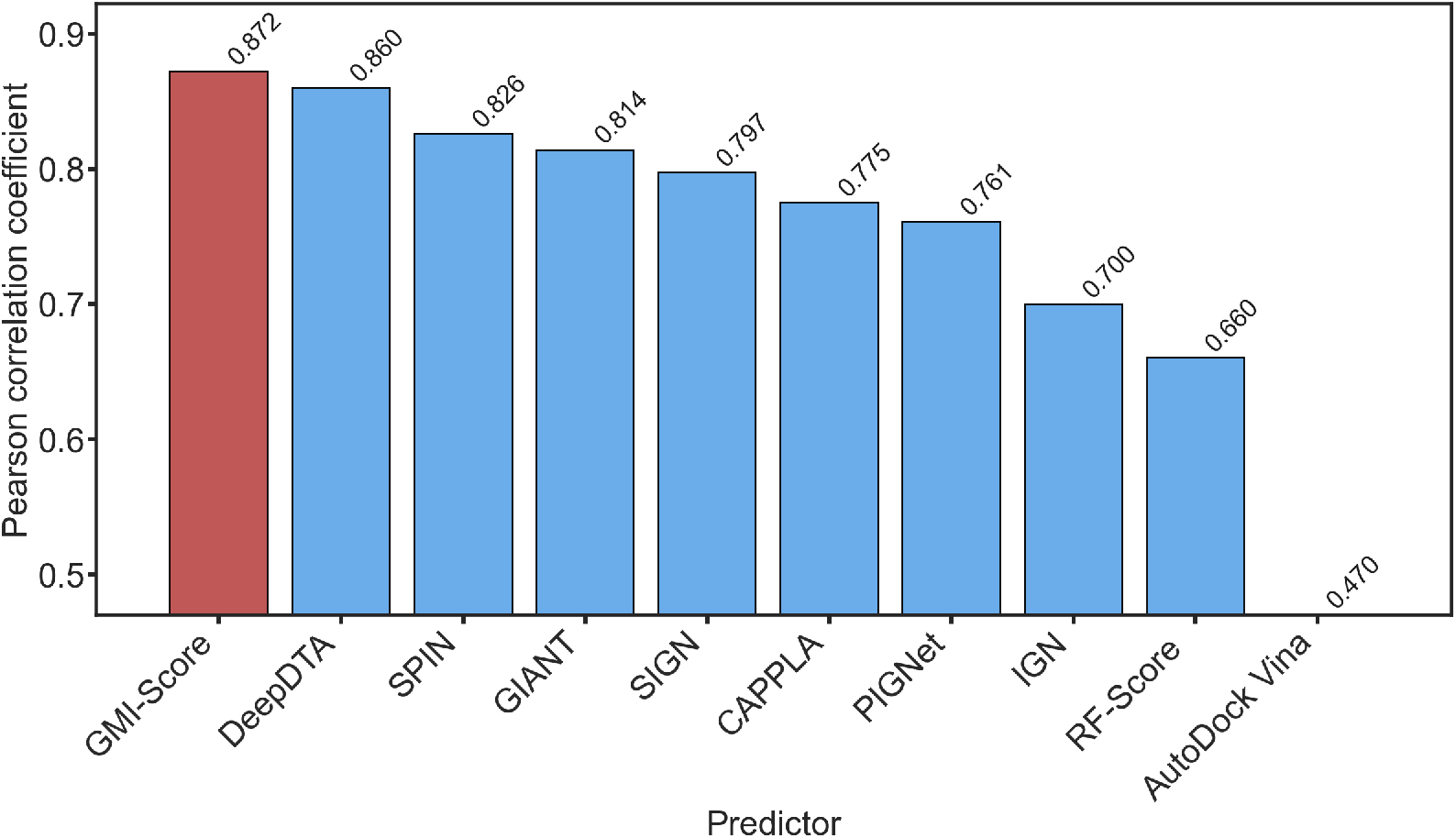
Comparison of Pearson’s correlation of various scoring functions trained on the PDBbind v2020 general set [4, 11, 25, 27, 33–35]. Our proposed *GMI-Score* is shown in red, while other methods are shown in blue. The performances of other models are taken from [11]. Only models achieving high *R*_*p*_ are reported.

### 2.5 PDBbind v2013 Benchmark

We further evaluated the generalization ability of GMI-Score through a cross-dataset validation strategy.The model was trained separately on the PDBbind v2020 general set and the PDBbind v2013 refined set, and then tested on the CASF-2013 core set. This benchmark comprises 195 high-quality protein-ligand complexes that are structurally diverse and span a broad range of binding affinities. Since none of these complexes overlap with the training sets, the CASF-2013 set provides an independent and unbiased assessment of model performance. The comparative results are presented in Figure 3.

**Fig. 3:**
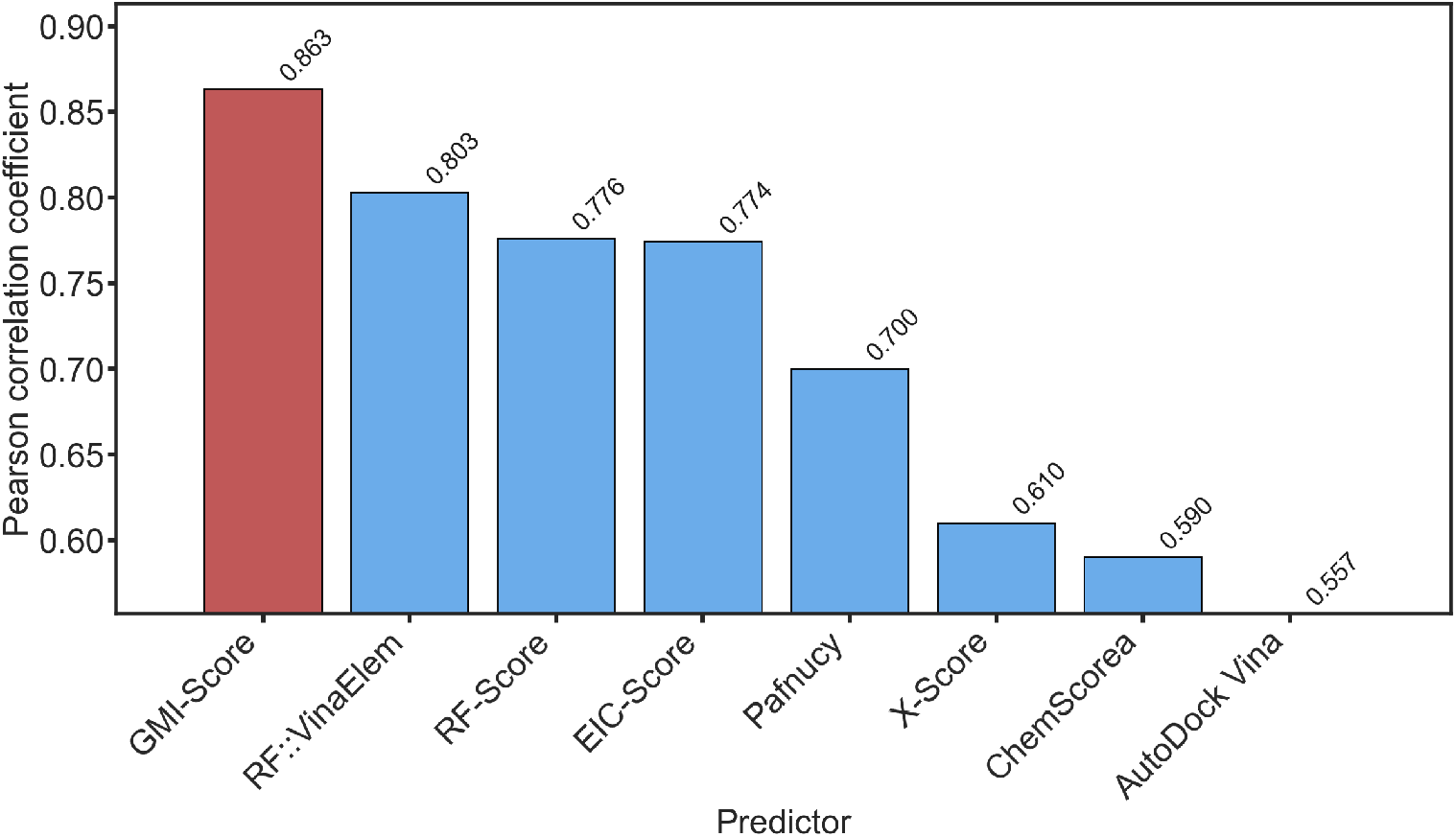
Comparison of Pearson’s correlation scores of various scoring functions of CASF-2013 set. Our proposed *GMI-Score* is shown in red, while other methods are shown in blue. The performances of other models are taken from [32, 43, 64, 72].

The GMI-Score achieves strong predictive performance on the CASF-2013 benchmark. Our method yields a higher correlation of 0.863 ± 0.001. This result demonstrates that GMI-Score maintains consistent accuracy across datasets of different sizes and compositions, which indicates its robust prediction to unseen protein-ligand systems.

### 2.6 CSAR-NRC-HiQ dataset

To further assess the generalizability of our GMI-Score, we conducted an external validation using a curated subset of the CSAR-HiQ dataset [13]. This dataset was filtered to remove any complexes that overlap with the PDBbind v2020 dataset (both general, refined and core sets), ensuring that our test set was truly independent. The final set comprised 49 unique complexes, which served as our independent validation set to evaluate the predictive power of the proposed model.

Predictions on the CSAR-NRC-HiQ dataset with the model trained on the PDB-bind v2020 general set show excellent performance. Our model achieved a Pearson’s correlation coefficient of 0.796 ± 0.016, with an associated *RMSE* of 2.321 ± 0.036 kcal/mol. It is worth noting that while GMI-Score achieves the highest *R*_*p*_ among all compared models, its *RMSE* (2.321 kcal/mol) is slightly higher than that of EGNA (2.094 kcal/mol). This seemingly contradictory result can be explained by the fact that *R*_*p*_ and *RMSE* capture complementary aspects of model performance. *R*_*p*_ reflects the model’s ability to correctly rank the relative binding affinities across complexes, whereas *RMSE* quantifies the absolute magnitude of prediction errors. A model can achieve strong ranking performance while exhibiting a degree of systematic offset or scale difference in its absolute predictions, particularly when optimization is targeted toward *R*_*p*_ as the primary metric. Furthermore, given the small size of the CSAR-NRC-HiQ set (49 complexes), *RMSE* values are more susceptible to the influence of individual outlier complexes. Taken together, the higher of GMI-Score suggests that it may provide a more reliable ordering of ligand binding strength, which is often the more practically relevant criterion in early-stage drug discovery workflows.

### 2.7 PDBbind-Redocked

To further assess the robustness of our framework under realistic drug-design conditions, we used the PDBbind-Redocked dataset [54] for this study. Unlike crystal-structure-based evaluations, this dataset was created by re-docking ligands from the PDBbind v2016 refined and core sets with Surflex-Dock [24], which generated up to 20 poses for each complex and selected the pose with the lowest RMSD among the top-scoring candidates. The redocking process imitates practical virtual-screening situations where experimental 3D structures are unavailable, providing a stringent and realistic test of a scoring function’s ability to generalize beyond idealized crystal data.

In this setup, the refined set (3,767 complexes) served as the training data, while the core set (285 complexes) was reserved as an independent test set. As shown in Table 5, GMI-Score reached the highest Pearson correlation coefficient (*R*_*p*_ = 0.829) among all compared models and shows strong consistency with experimental binding affinities. GatedGCN 1.0 had slightly a lower correlation (*R*_*p*_ = 0.810). The relatively small gap between GMI-Score and GatedGCN 1.0 suggests that both models capture meaningful interaction patterns even from computationally generated poses. AutoDock Vina showed the weakest performance (*R*_*p*_ = 0.521). The considerably weaker performance of AutoDock Vina is consistent with its well-documented limitations in absolute affinity estimation [62], and aligns with findings from other external validation studies showing that physics-based scoring functions tend to struggle more than machine learning-based approaches when applied to non-native conformations [59, 70]. Overall, the high correlation of GMI-Score demonstrates that it provides a more reliable rank order of ligand binding strength, even for docking-generated conformations. This result highlights the model’s potential in real-world drug-design applications where crystal structures are often unavailable.

**Table 5:**
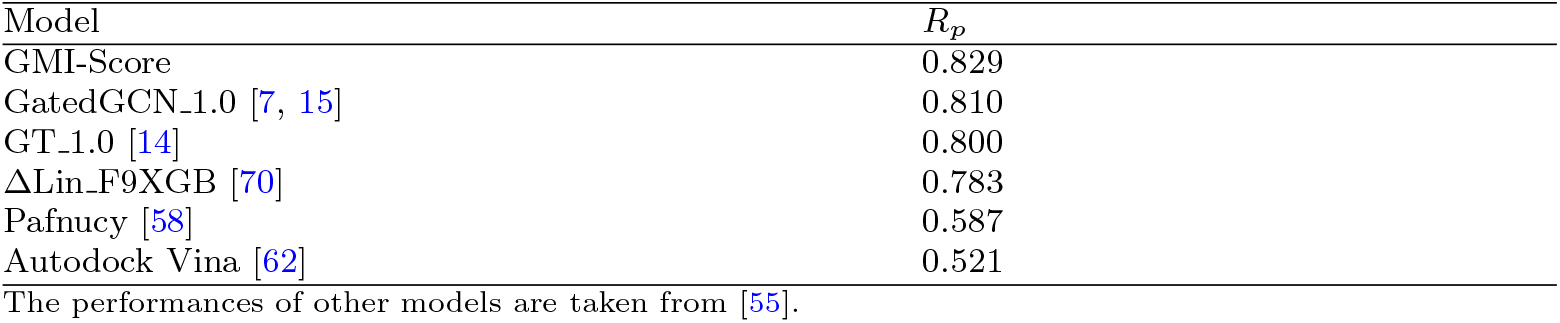
Performance comparison on the PDBbind-Redocked dataset.

| Model | $R_p$ |
| --- | --- |
| GMI-Score | 0.829 |
| GatedGCN_1.0 [7, 15] | 0.810 |
| GT_1.0 [14] | 0.800 |
| ΔLin_F9XGB [70] | 0.783 |
| Pafnucy [58] | 0.587 |
| Autodock Vina [62] | 0.521 |

The performances of other models are taken from [55].

## 3 Methods

### 3.1 Multi-geometric subgraphs

Understanding the complex interactions within biomolecules often begins by representing them as graphs. Existing works have experimented on a variety of schemes to classify the atomic interactions. Early approaches [41, 44] classified the interactions based on element symbols of the protein-ligand atoms, leading to coarse interaction types such as C-N or C-O. In more recent work [50], two additional schemes have been utilized to classify atomic interactions in protein-ligand complexes. In the GGL-Score scheme[50], protein atoms (excluding hydrogens) are identified by their atom names, and ligand atoms are described using SYBYL atom types, resulting in more detailed interaction pairs such as CD1-C.2, CG-C.ar, or the second scheme considers interaction between heavy atoms of each complex using a geometric subgraph of many clusters of pairwise and triplet-wise bonding space. Let 𝒯 be the set of all interested atom types of any of the schemes above.

**Fig. 4:**
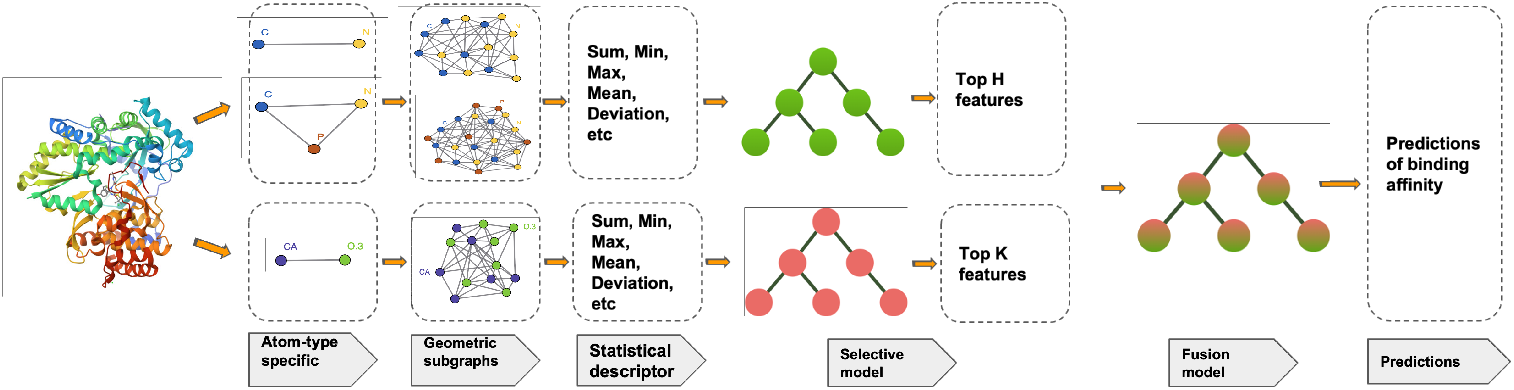
Illustration of GMI strategy using 2GNW complex. In the second and third columns, lement-specific clusters of GMI-Score-3 and GMI-Score-2 are presented, respectively, from top to bottom. The statistical descriptors (fourth column) are selected by a feature selection method using a machine learning model, then a fusion technique is applied before training and predicting with a gradient boosting trees model.

**Fig. 5:**
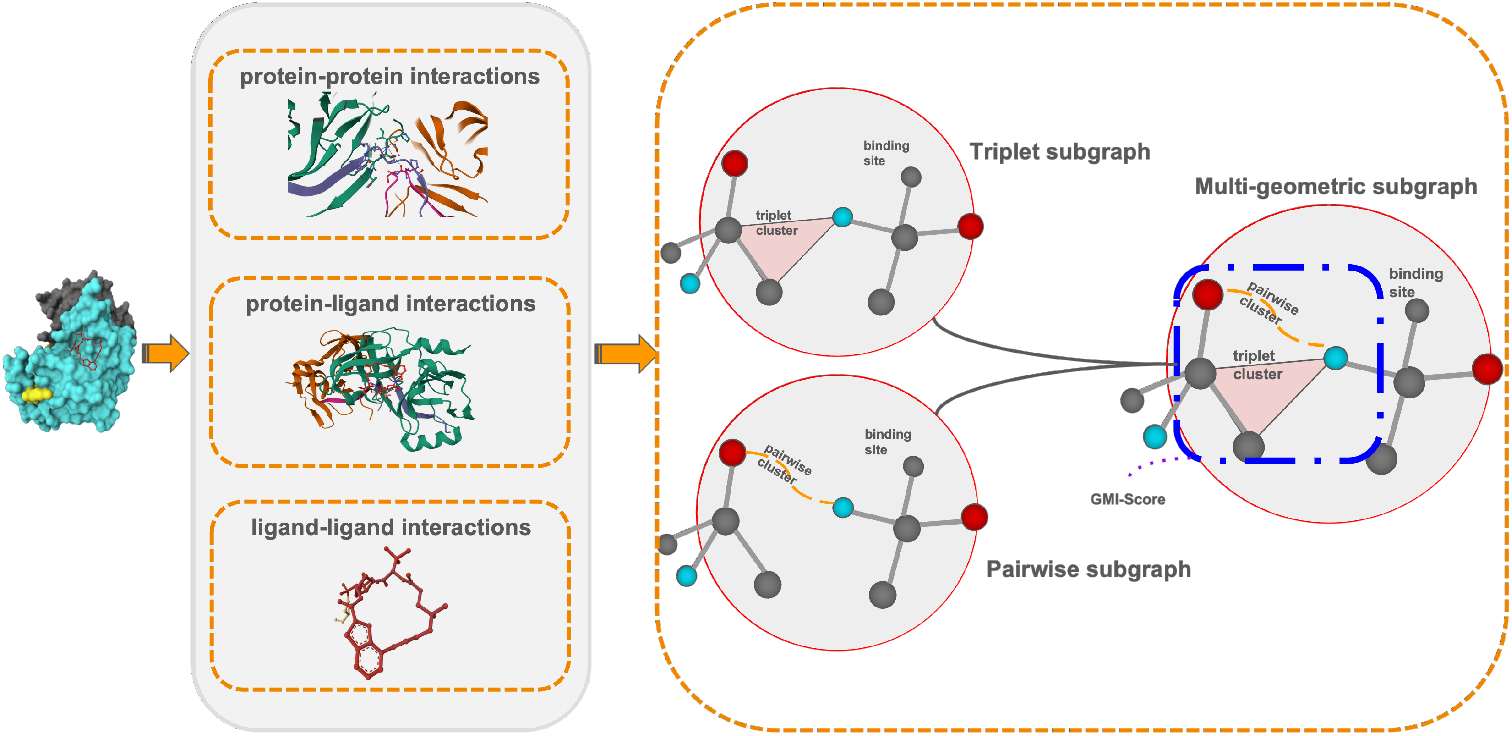
Multi-geometrics subgraphs (multibody subgraphs) are the combination of triplet subgraphs and pairwise subgraphs.

#### 3.1.1 Complete weighted molecular graph

We represent a protein-ligand complex as a complete weighted molecular graph 𝒢= (𝒱, *ε, w*). Each vertex corresponds to an atom in the complex. Let the entity labels be = {P, L}, denoting protein (P) and ligand (L) atoms. Each entity is associated with its own set of chemical atom types

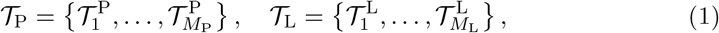

where *M*_P_ and *M*_L_ denote the numbers of distinct atom types in the protein and ligand, respectively. The augmented type set is the disjoint union

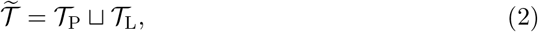

so that each atom is uniquely characterized by both its chemical type and its entity label.

The vertex set is

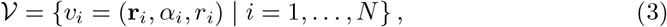

where *N* is the number of considered atoms in the protein-ligand complex, **r**_*i*_ ∈ ℝ^3^ is the coordinate of atom 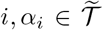 is its augmented type, and *r*_*i*_ is its van der Waals radius. The edge set includes all unordered pairs of distinct atoms:

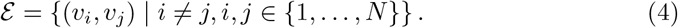

Thus, the molecular graph is complete, and every atom pair is connected. Each edge (*v*_*i*_, *v*_*j*_) is assigned a type-dependent weight

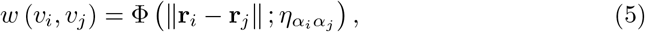

where Φ is a kernel function parameterized by a characteristic distance 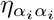, we place the following conditions on the weight function Φ as previous work [44]

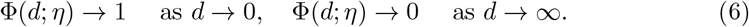

Evidence from previous works [44, 50] have shown that the generalized exponential function:

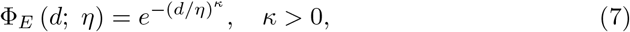

and the generalized Lorentz function

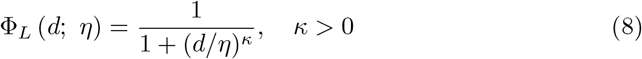

are good candidates for the non-covalent interactions.

This complete weighted molecular graph captures the full atomistic structure of the protein-ligand complex and serves as the foundation for defining type-specific subgraphs and multi-body interaction scores.

#### 3.2.1 Multi-body type-specific subgraphs

While the complete weighted molecular graph 𝒢 captures all pairwise interactions in the protein-ligand complex, practical analysis often requires focusing on subsets of atom types to isolate physically meaningful multi-body interactions.

Let

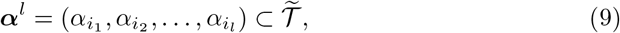

denote an ordered collection of *l* augmented atom types drawn from the combined set 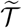. The vertex set associated with ***α***^*l*^ is

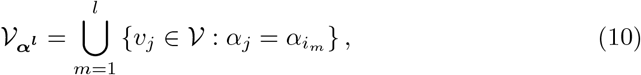

and the edge set is

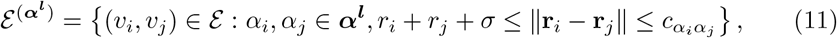

where 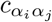 is a type-pair-specific cutoff distance. This parameter restricts edges to atom pairs that are spatially close enough to form chemically relevant interactions. The cutoff may be chosen as a fixed constant for each type pair or adaptively defined from empirical distributions of interaction distances. *σ* is the mean value of the standard deviation of the radii in the dataset.

If one is interested in considering only the intermolecular interactions between protein and ligand, the edge set will be defined as follows

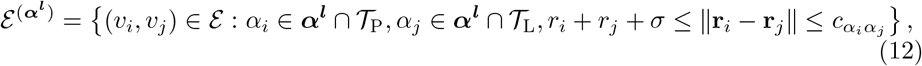

The resulting multi-body type-specific subgraph for *l* atom types is

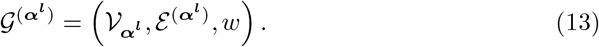

This framework accommodates arbitrary selections of protein and ligand atom types, giving rise to the construction of subgraphs that capture intermolecular interactions (protein-ligand), intramolecular interactions (protein-protein, ligand-ligand), as well as higher-order cooperative subgraphs involving multiple atom types simultaneously. When *l* = 2, we obtain the pairwise atom type-based models as mentioned in our previous work [44, 50].

#### 3.1.3 Multi-body interaction scores

The construction of multi-body type-specific subgraphs provides a natural framework for quantifying cooperative interactions among selected atom types in the proteinligand complex. While pairwise kernels *w* (*v*_*i*_, *v*_*j*_) capture local atomic affinities, they describe only isolated atom–atom contacts and do not scale naturally to reflect the collective behavior of larger subgraphs. Thus, we propose multi-body interaction scores, which aggregate edge weights across *l*-atom types subgraphs to measure the strength of cooperative interactions within a chosen set of atom types. We have two different kinds of the multi-body (MB) scores:

##### Cyclic interaction Score

For an ordered *l*-tuple 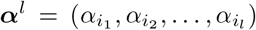 with cyclic closure 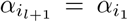, the cyclic score is

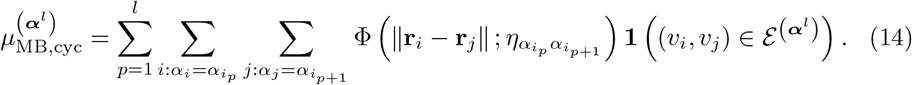

##### Clique interaction score

Alternatively, the clique score aggregates all pairwise couplings among the chosen types

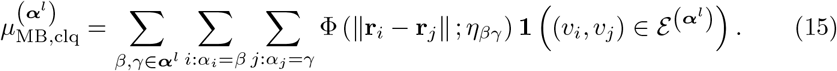

The operator 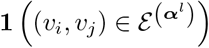 is an indicator function

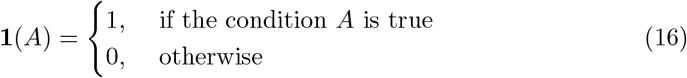

Since this work does not consider the ordering of atom types, only the clique interaction score is employed in all experiments. Moreover, to avoid redundant feature representations, we restrict attention to the pairwise multi-body model (GMI-Score-2) with *l* = 2, and the triplet multi-body model (GMI-Score-3) with *l* = 3.

### 3.2 Fusion learning

Many existing works demonstrate that combining various descriptors can enhance the prediction of protein-ligand binding affinities [28, 31, 65, 69]. Moreover, the importance of data quality over quantity, specifically, the relevance and informativeness of descriptors are shown to have a substantial impact on model performance and efficiency [41]. To obtain richer representations of the protein-ligand omplexes, we fuse the most informative features from both GMI-Score-2 and GMI-Score-3. Particularly, we focus on subsets of the pair and triplet atom types from both feature sets, which not only preserves key biochemical interactions but also eliminates non-contributory features, thereby reducing the computational cost. GMI-Score-2 provides the interaction strengths between atom pair types, whereas GMI-Score-3 captures both pairwise and triplet-wise energy terms across multi-body interactions, together forming multi-level representations of protein-ligand interactions.

To capitalize on the complementary strengths of both methods, we adopted a fusion strategy. The rationale behind combining different scoring functions or models is well-established in computational drug discovery [10]. Ensemble methods, such as those employing combinatorial fusion analysis [23], often lead to improved accuracy and robustness by mitigating the individual limitations and biases of single models [40]. Here, we developed a new SF model by integrating GMI-Score-2 and GL-MB-Score-3 via a fusion process. The process can be done through methods like weighted averaging, linear regression, or more sophisticated ensemble techniques such as stacking or boosting. Such methods focus on leveraging the individual predictive power of each model at the decision stage. In our work, we applied an ensemble learning strategy that integrates their outputs at the decision stage. Specifically, we adopted a rank-based aggregation method: ensemble models ranked all candidate ligand–protein complexes according to their predicted binding affinity score. The final ensemble ranking was then obtained by aggregating these individual ranks, such that the relative order of complexes reflected the consensus between the two models.

#### 3.2.1 Feature selection

Gradient boosting is a machine learning method that builds an ensemble of weak prediction models (typically decision trees) in a stage-wise fashion, optimizing an arbitrary differentiable loss function using gradient descent [18]. At each iteration, the algorithm fits a new model to the pseudo-residuals of the current ensemble, gradually reducing prediction errors [42]. Formally, given training data 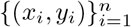 and a differentiable loss function *L*(*y, F* (*x*)), the method builds an additive model 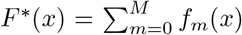 where each *f*_*m*_ is a weak learner that minimizes the expected loss [18].

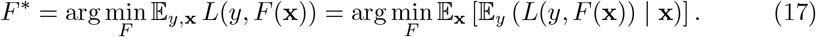

where 𝒢 (*y, F* (**x**)) is the loss function. Decision trees are developed using the greedy CART algorithm. During this process, features are chosen selectively to maintain sparsity in order to minimize the total impurity of child nodes [6]. Given a node *t* and its probabilities *p*(*j*|*t*), the impurity is calculated as follow:

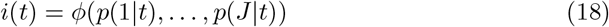

Gini index is then applied to rank the importance of variables can be written as:

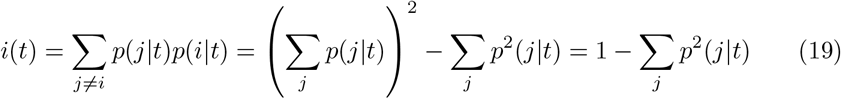

#### 3.2.2 Feature fusion

To effectively merge two distinct feature sets while ensuring we capture their most informative descriptors, we employed a strategic feature fusion technique. This method centers on assessing the importance of each feature, a widely recognized approach for pointing out influential predictors within data. Rather than simply combining or averaging features, which can lead to redundancy or dilute vital information, our process utilizes the feature ranking of Gradient Boosting (GB) models.

Given *n* minimum node impurity of features from *m* datasets 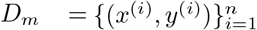, the new variables are defined as:

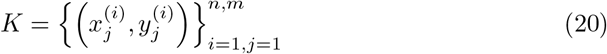

This concatenation creates a richer representation by combining the most informative descriptors from both original sets, allowing the subsequent model to leverage a broader spectrum of predictive signals.

#### 3.3 Benchmarck evaluation

In this work, our proposed GMI-Score was evaluated for scoring power across several datasets. Following the standard CASF benchmark protocol [59] for scoring power, the Pearson’s correlation coefficient (R p) between the predicted binding scores and the experimental binding constants (in logarithm units, log Ka) was computed as a quantitative indicator of scoring power:

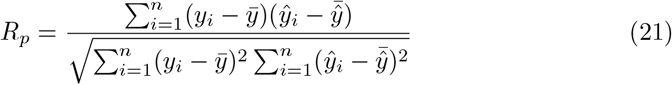

To account for variability, each experiment was repeated at least 5 times with different random seeds, and the reported results represent the average across runs. RMSE values are optionally included where available to facilitate comparison with other methods.

## 4 Conclusions

In this work, we introduced GMI-Score, a novel scoring function for protein-ligand binding affinity prediction that leverages multi-level, multi-body atomic interaction graphs. Unlike traditional approaches that depend heavily on handcrafted physico-chemical descriptors, our method utilizes graph-based representations as core features, significantly reducing model complexity while maintaining high predictive accuracy. By integrating both pairwise and triplet-wise interactions through a feature fusion strategy, GMI-Score effectively captures the physical and chemical information governing the protein-ligand binding mechanism. Our extensive experiments on multiple benchmark datasets, including PDBbind v2016, v2020, CSAR-NRC-HiQ, and PDBbind-Redocked, demonstrate consistent improvements over state-of-the-art methods. Our proposed models achieve Pearson correlations up to 0.877. Among all evaluated ML-based models, GMI-Score exhibits the smallest performance degradation on the leak-proof partition, further supporting its robustness on unseen and diverse molecular systems.

Despite these promising results, this study focuses solely on the scoring power measured by the Pearson correlation coefficient following the PDBbind protocol. Other important evaluation criteria, including ranking power, docking power, and screening power, which require additional dataset preparation and methodological extensions, were not assessed and remain the main direction for future work.

## Author contributions

T.TH.L, H.V, N.HM.N prepared the manuscript; T.TH.L, B.T.N, D.D.N finalized the manuscript. T.TH.Lcollected data and performed analysis. B.T.N, D.D.N supervised the project.

## Funding

D.D.N. was supported in part by funds from the National Science Foundation (NSF: # 2516126, # 2151802, and # 2534947).

## Data availability

The PDBbind data used in this study are publicly available at https://www.PDBbind-plus.org.cn. The source code and pretrained model are available at the GitHub repository: https://github.com/BinhMisfit/DrugWise-Implementation.

## Declarations

### Competing interests

The authors declare that they have no competing interests.

## Appendix A Supplementary Tables

**Table A1:** Model hyperparameters including number of estimators, feature selection method, tree depth,split criteria, learning rate, subsampling ratio, and loss function.

| **Model** | **Hyperparameters** |
| --- | --- |
| Model 1 | n_estimators=20000, max_features='sqrt', max_depth=8, min_samples_split=2, learning_rate=0.005, subsample=0.7, loss='squared_error' |
| Model 2 | n_estimators=15000, max_features='sqrt', max_depth=7, min_samples_split=16, min_samples_leaf=10, learning_rate=0.008, subsample=0.4, loss='squared_error' |
| Model 3 | n_estimators=24500, max_features='sqrt', max_depth=7, min_samples_split=3, min_samples_leaf=19, learning_rate=0.03, subsample=0.4, loss='squared_error' |
| Model 4 | n_estimators=23500, max_features='sqrt', max_depth=6, min_samples_split=16, min_samples_leaf=8, learning_rate=0.006, subsample=0.7, loss='squared_error' |

**Table A2:** Paired t-test on Pearson correlation: GMI-Score vs individual model on CASF v2016 dataset.

| Model | Train with Refined |  |  | Train with General |  |  |
| --- | --- | --- | --- | --- | --- | --- |
|  | t-stat | p-value | significant | t-stat | p-value | significance |
| GMI-Score-2 | 16.163 | 0.0001 | *** | 13.191 | 0.0002 | *** |
| GMI-Score-3 | 10.090 | 0.0005 | *** | 25.142 | 0.0000 | *** |

Significance: \*\*\*: $p < 0.001$ | \*\*: $p < 0.01$ | \*: $p < 0.05$ | n.s. = not significant.

**Table A3:** Performance comparison of individual and fusion models on the PDBbind v2020 dataset. The table reports the Pearson correlation coefficient (*R*_*p*_) and RMSE in kcal/mol.

| Model | Train with Refined |  | Train with General |  |
| --- | --- | --- | --- | --- |
| | $R_p$ | $RMSE$ | $R_p$ | $RMSE$ |
| GMI-Score-2 | $0.816 \pm 0.001$ | $1.829 \pm 0.003$ | $0.862 \pm 0.002$ | $1.607 \pm 0.008$ |
| GMI-Score-3 | $0.834 \pm 0.002$ | $1.769 \pm 0.009$ | $0.863 \pm 0.001$ | $1.622 \pm 0.007$ |
| GMI-Score | $0.842 \pm 0.002$ | $1.732 \pm 0.008$ | $0.872 \pm 0.001$ | $1.560 \pm 0.003$ |

Significance: \*\*\*: $p < 0.001$ | \*\*: $p < 0.01$ | \*: $p < 0.05$ | n.s. = not significant.

**Table B4:** Paired t-test on Pearson correlation: GMI-Score vs individual model on PDBbind v2020 dataset.

| Model | Train with Refined |  |  | Train with General |  |  |
| --- | --- | --- | --- | --- | --- | --- |
|  | t-stat | p-value | significant | t-stat | p-value | significance |
| GMI-Score-2 | 23.791 | 0.0000 | *** | 13.870 | 0.0002 | *** |
| GMI-Score-3 | 5.396 | 0.0057 | *** | 13.425 | 0.0002 | *** |

Significance: \*\*\*: $p < 0.001$ | \*\*: $p < 0.01$ | \*: $p < 0.05$ | n.s. = not significant.

## Appendix B Supplementary Figures

**Fig. B1.**
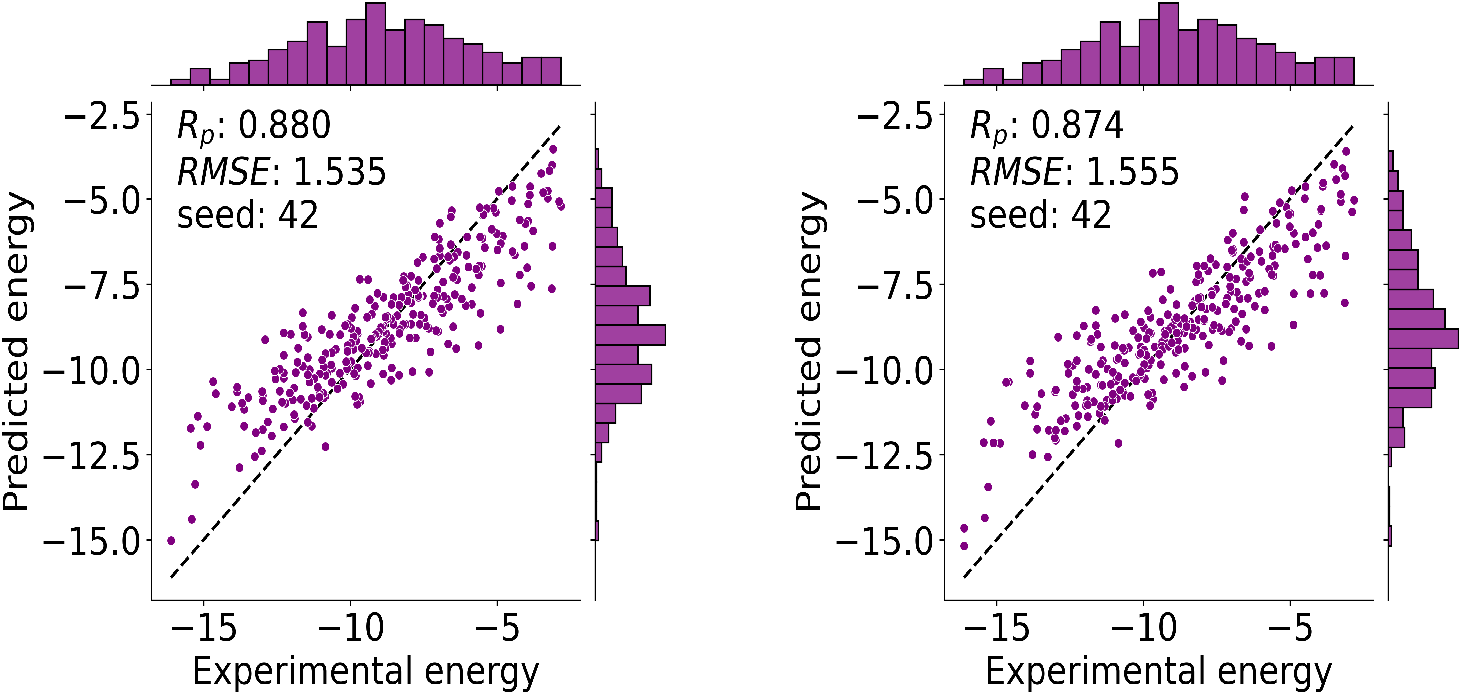
Correlation between GMI-Score predictions and experimental data for various benchmarks. (Left) PDBbind v2016: Pearson correlation coefficient *R*_*p*_ = 0.880 and RMSE = 1.535 kcal/mol. (Right) PDBbind v2020: Pearson correlation coefficient *R*_*p*_ = 0.874 and *RMSE* = 1.555 kcal/mol.

## Appendix C Predictions of PDBbind v2016 dataset

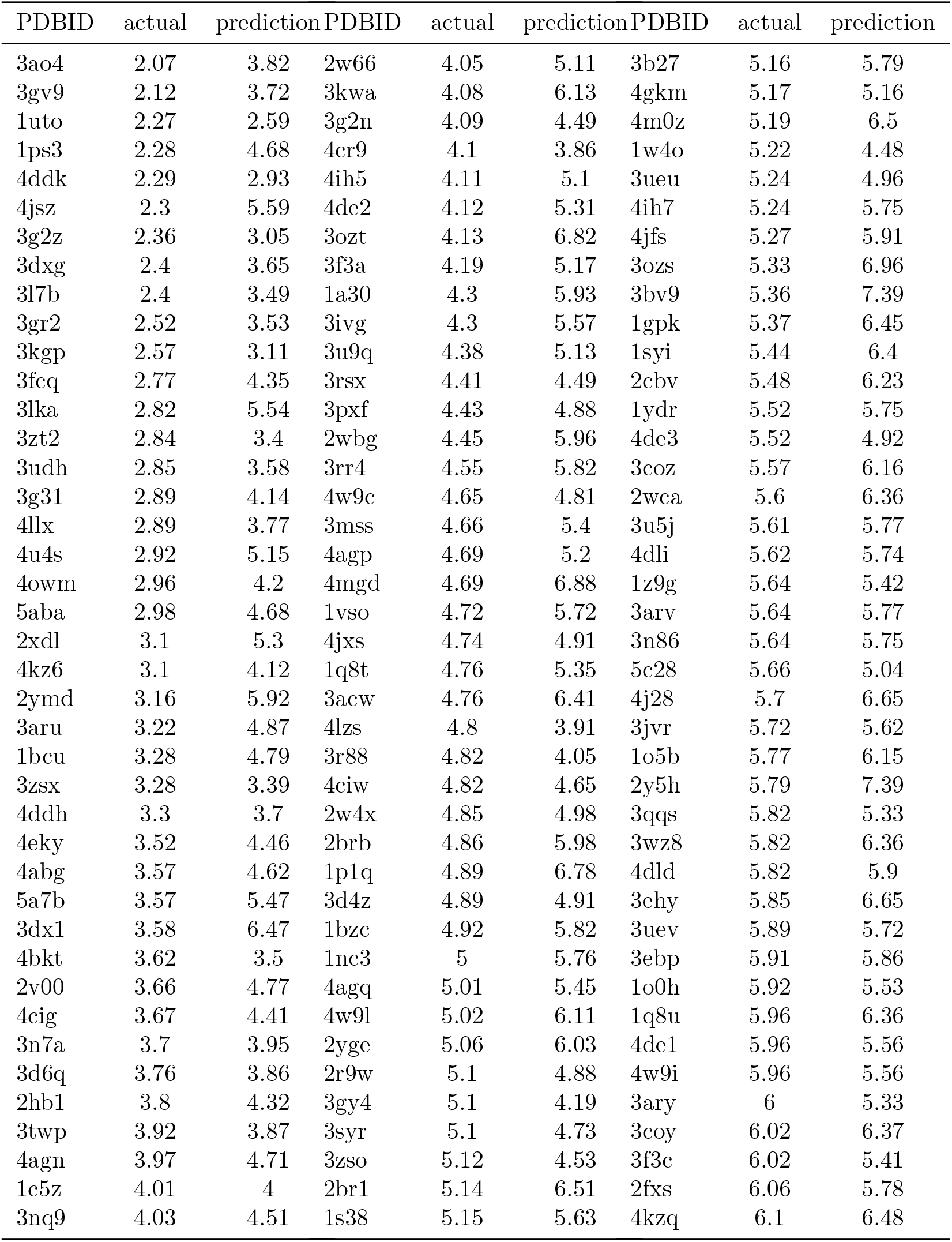
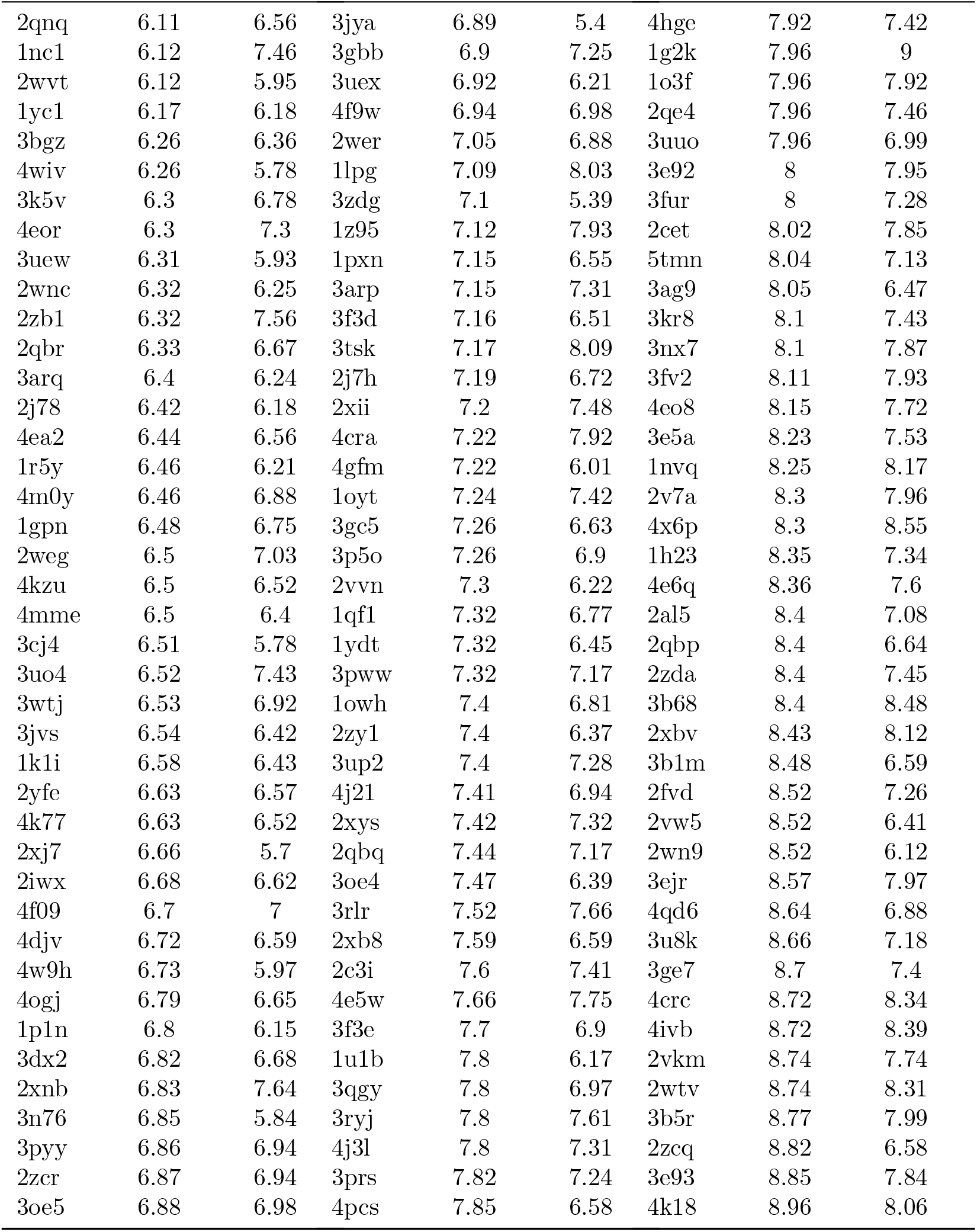
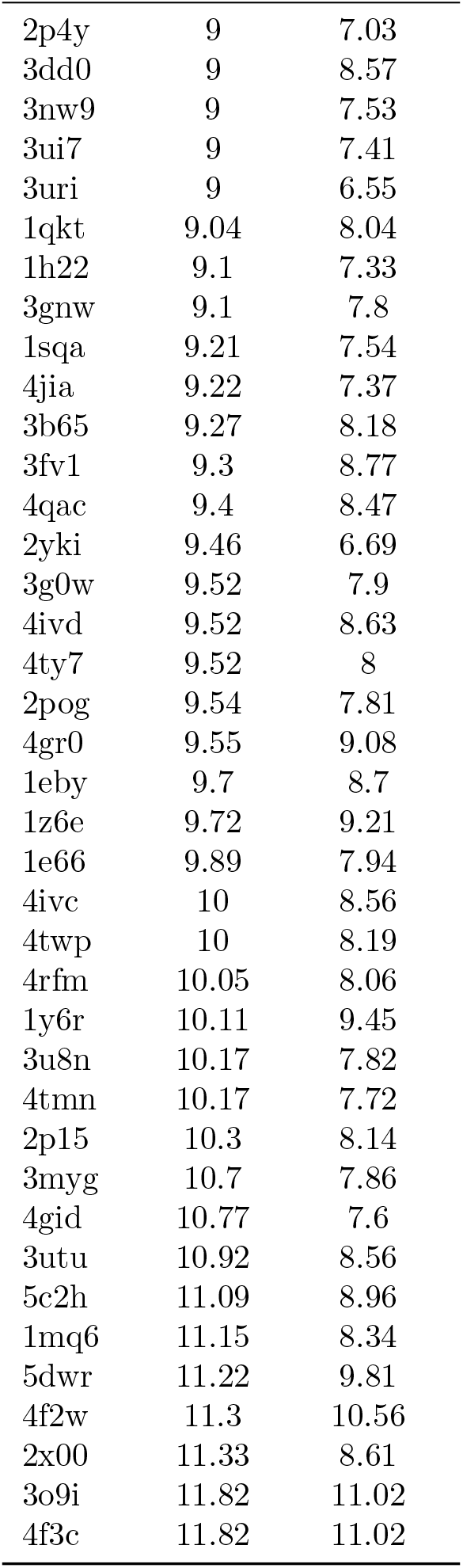

## Appendix D Predictions of PDBbind v2020 dataset

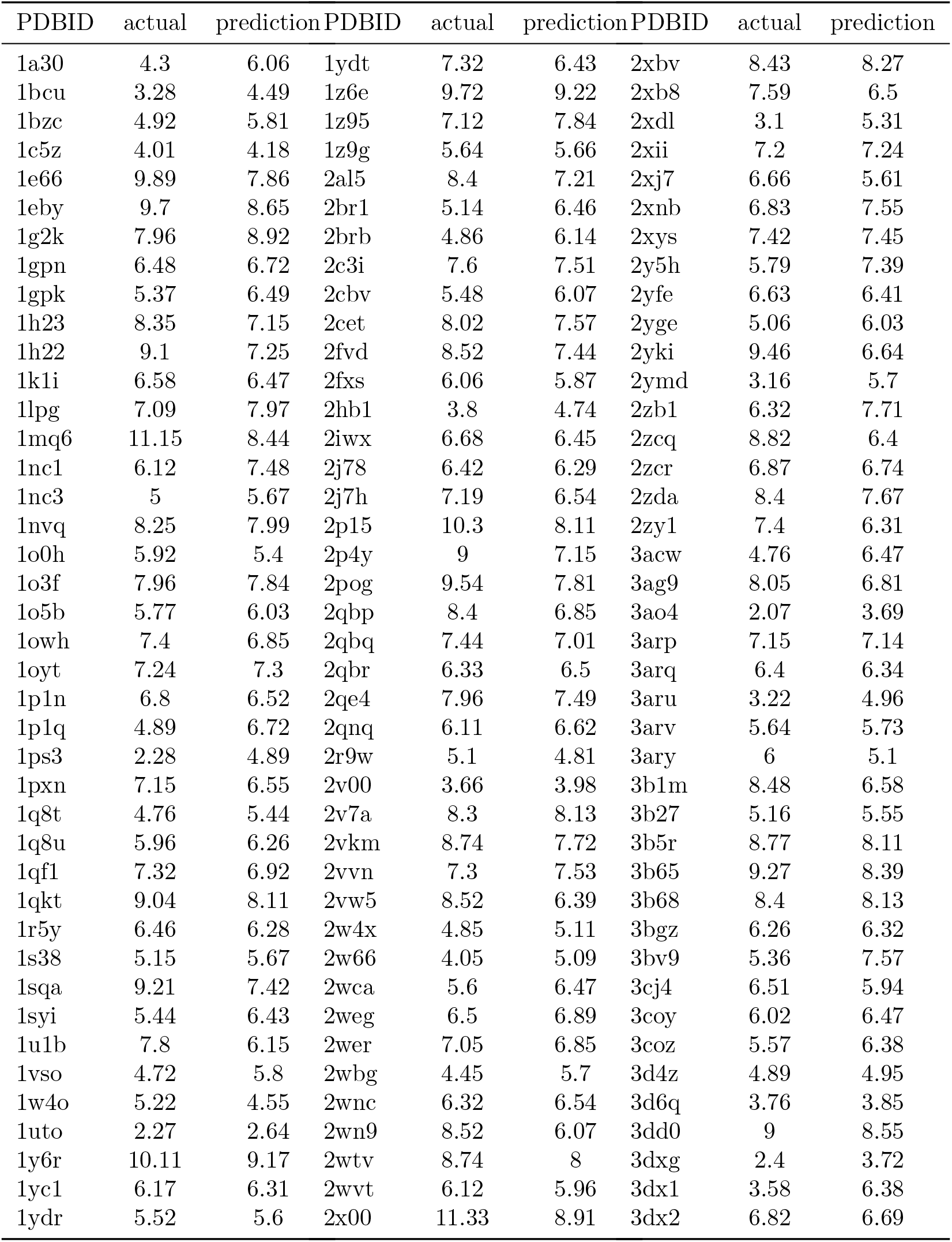
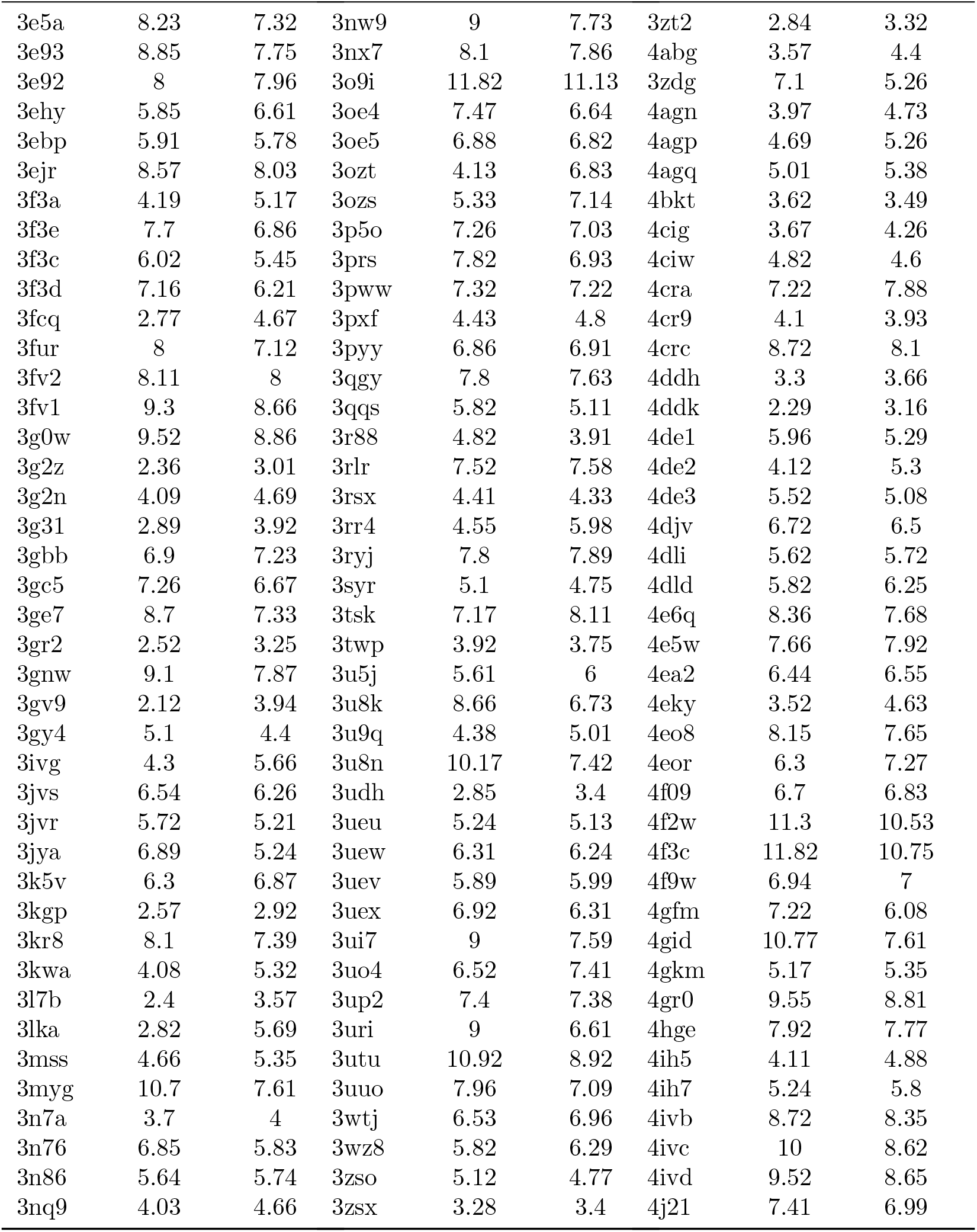
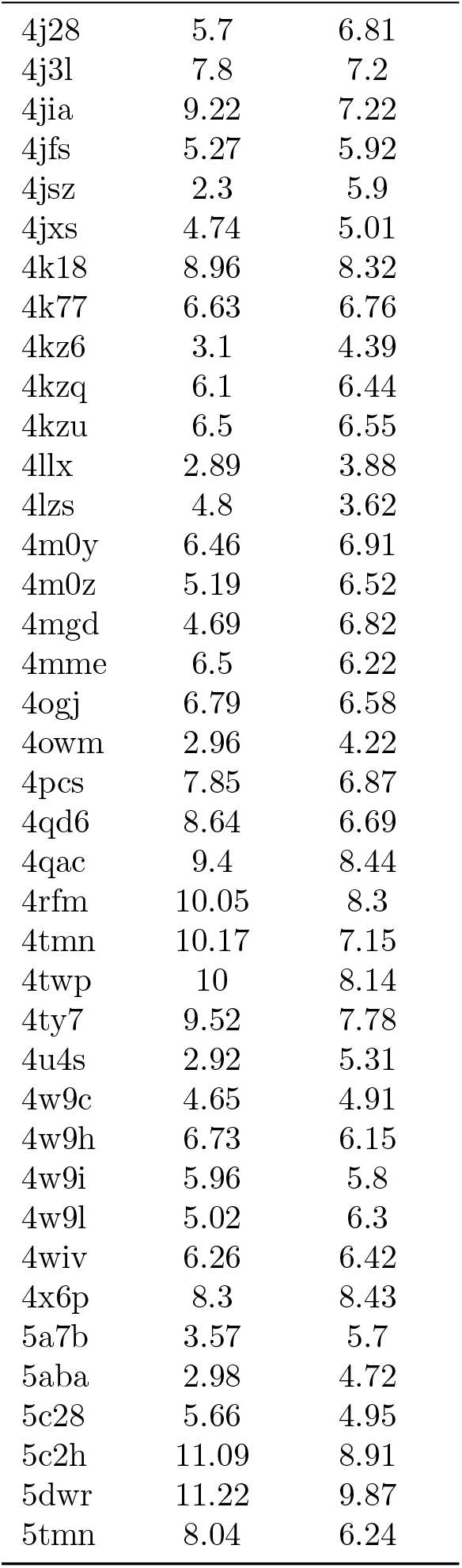

## Notes

https://www.PDBbind-plus.org.cn

### Competing Interest Statement

The authors have declared no competing interest.

### Summary of Updates

Correct the author's name (change 'Tran' to 'Tram').

